# Estimating seven coefficients of pairwise relatedness using population genomic data

**DOI:** 10.1101/049411

**Authors:** Matthew S Ackerman, Parul Johri, Ken Spitze, Sen Xu, Thomas Doak, Kimberly Young, Michael Lynch

## Abstract

Population structure can be described by genotypic correlation coefficients between groups of individuals, the most basic of which are the pair-wise relatedness coefficients between any two individuals. There are nine pair-wise relatedness coefficients in the most general model, and we show that these can be reduced to seven coefficients for biallelic loci. Although all nine coefficients can be estimated from pedigrees, six coefficients have been beyond empirical reach. We provide a numerical optimization procedure that estimates them from population-genomic data. Simulations show that the procedure is nearly unbiased, even at 3× coverage, and errors in five of the seven coefficients are statistically uncorrelated. The remaining two coefficients have a negative correlation of errors, but their sum provides an unbiased assessment of the overall correlation of heterozygosity between two individuals. Application of these new methods to four populations of the freshwater crustacean *Daphnia pulex* reveal the occurrence of half-siblings in our samples, as well as a number of identical individuals that are likely obligately asexual clone-mates. Statistically significant negative estimates of these pair-wise relatedness coefficients, including inbreeding coefficents that were typically negative, underscore the difficulties that arise when interpreting genotypic correlations as estimations of the probability that alleles are identical by descent.

## Introduction

While some phenotypic variation within a population results from variation in single genes, many phenotypes are influenced by complex interactions between multiple genes and various environmental conditions (Fisher 1918). It may be impossible to isolate all of the genetic factors contributing to a complex phenotype, but the total genetic contribution to phenotypic variation can be estimated using regression coefficients that describe the statistical association between the genotypes of two individuals.

This statistical association is usually attributed to the sharing of haplotypes, or individual copies of genes, that are identical by descent (IBD), which occurs when two or more haplotypes descend from a single ancestral haplotype (Epperson 1999). When considering groups of more than two haplotypes, a number of IBD coefficients are necessary to describe the probability that haplotypes within each subset of the group are IBD. If the genotype to phenotype relationship were very simple, then a single measure of genotypic correlation which described how the number of alleles are correlated between individuals—the coefficient of coancestry (Θ)—would be sufficient to relate genetic covari-ance to pheotypic covariance. Unfortunately, many genes do not follow a simple additive model of gene action, and as a result additional genotypic correlation coefficients are needed.

For example, siblings tend to have a stronger phenotypic resemblance to each other than either sibling has to their parents. While this may seem surprising, it can be understood as a result of non-additive gene action. If the parents are unrelated to each other, then at every locus a single parent and offspring share exactly 1 pair of alleles that are IBD (Θ = 0.25, see Table S5). However, heterozygosity and homozygosity (jointly called zygosity) are defined by the relationship of two haploid genomes to each other. Because each parent gives only one haploid genome to their offspring, the offspring’s zygosity is unassociated with the zygosity of their parent, so the coefficient of cofraternity (Δ)— which describes the association of zygosity between individuals—is zero. While siblings share alleles that are identical by descent with each other with equal probability that siblings share alleles that are identical by descent with their parents, siblings can receive the same alleles from both their parents, creating a correlation of zygosity state (Δ ≈ 0.25) in addition to the correlation of allele count. Because alleles often exhibit some form of dominance, a pair of siblings generally has greater genetic covariance than a parent-offspring pair, despite having similar coancestry coefficients (Lynch and Walsh 1998).

In a randomly mating, outbred population, knowledge of both Θ and Δ between all individuals is sufficient to relate the genetic covariance of individuals, *σ*_G_(*X,Y*), to the additive (*σ*_A_^2^) and dominance (*σ*_D_^2^) genetic variation in a population (Lynch and Walsh 1998). These terms are used to estimate heritability, and thus IBD coefficients are fundamental to a variety of quantitative-genetic analyses. However, Θ and Δ are only sufficient to estimate genetic variation in panmictic outbred populations. Additional genetic variance terms and IBD coefficients are necessary to describe inbred individuals.

General genotypic correlation coefficients can be estimated if the pedigree of the related individuals is known (Wright 1922). However, because each round of reproduction involves a limited number of crossover events, typically on the order of one event per chromosome arm, the actual pattern of inheritance can vary substantially from the expectation predicted by path analysis. For instance, a pair of human half siblings have an expected coancestry of Θ = 0.125 but since ~184 ^1^ crossover events separate them, ~5%^2^ of half siblings will have a coancestry that is less than 0.092 or greater than 0.158 (Speed and Balding 2015).

Some of the shortcomings of pedigree analysis can be addressed by estimating genotypic correlation coefficients from molecular markers (Lynch and Ritland 1999, Wang 2002, Fernández and Toro 2006, Kalinowski et al. 2006, Wang 2007, Anderson and Weir 2007, Wang 2011). However, different conceptualizations of genotypic correlation have resulted in variety of estimators that use a range of analytical techniques. Genotypic correlation is frequently defined as representing the probability that two sequences descended from a single ancestral sequence without recombination. This recombination-based definition focuses statistical analysis on the distribution of lengths of IBD tracks (Sved 1971). Additionally, defining genotypic correlations as the probability of IBD limits meaningful estimates to the interval between 0 and 1. While neither of these properties are undesirable *per se*, using these coefficients as regression coefficients is undesirable because it creates bias in estimates of heritablity (Lynch and Ritland 1999).

Alternatively, genotypic correlation coefficients can be defined as the coefficients that describe the genetic covariance of quantitative traits. This approach has been taken for a number of marker-based methods, but these methods make restrictive assumptions about the pedigrees of individuals, excluding the possibility of inbreeding or outbreeding (Lynch and Ritland 1999, Wang 2007); or they require a large number of alleles at a locus to achieve substantial statistical power (Fernández and Toro 2006 or Wang 2011), while the most abundant markers in modern data sets are biallelic SNPs (representing only two alleles).

With these problems in mind, we sought to develop a method for estimating relatedness that makes effective use of the bial-lelic markers abundantly available in population-genomic data, without making restrictive assumptions about possible values of relatedness coefficients. We show how both genotypic and phenotypic correlation coefficients can be negative, which emphasizes that these coefficients are not probabilities, and also show that seven coefficients, rather than nine, are sufficient to specify the genetic covariance at biallelic loci.

## Genotypic correlation

### A statistical view of genealogies

The genotype of one individual often gives us some information about the genotypes of other individuals. For instance, we expect the members of a species to be genetically similar, so sequencing a single individual of that species can give us some idea of the genes present in most members of that species. This genetic similarity arises in part from the common ancestry of all members of a species. The metaphor of identity by descent captures this part of the explanation, but also obscures the influence of mutation and genetic drift on these correlations. This lacuna of understanding is also present in pedigree-based calculations of IBD coefficients, which, despite over a hundred years of use, can only be calculated from truncated pedigrees. Calculations from exhaustive pedigrees cause pedigree-based calculations of genotypic correlations to approach one (Speed and Balding 2015), differing sharply from the IBD coefficients calculated using a molecular-marker method. A careful consideration of the statistical processes at work highlights the roles of mutation and drift in generating genotypic correlations, and shows how molecular-marker method are related to pedigree-based methods.

We can imagine that an individual’s genotype is determined by a three step process, in which an allele (SNP) originates in some particular ancestor through mutation (individual *Z* in Figure 1), then descends stochastically down a fixed genealogical structure, and ultimately comes to rest in the genome of the individual that we have sampled (individual *X* or *Y* in Figure 1). The genotypic correlation between two gametes is a measure of the tendency of alleles to co-occur in those gametes, and in order to calculate this correlation we will need to estimate three probabilities—the probability of sampling a particular allele in gamete *a*, *P*(*a*), the probability of sampling that allele in gamete *b*, *P*(*b*), and the probability of sampling the allele in both gametes simultaneously, *P*(*ab*) (see Table 1 for other symbols and their definitions).

**Figure 1.**
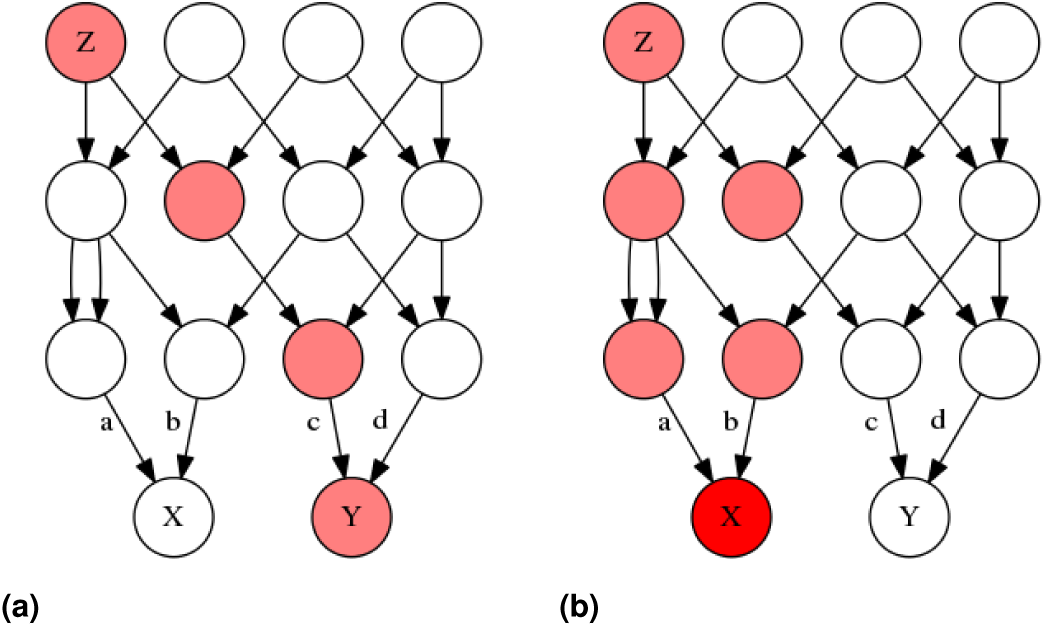
Two genealogies of identical structure which illustrate the stochastic model of genotypic correlations. In both of these genealogies a mutation occurs in ancestor *Z*, making *Z* heterozygous for some trait (pink). The allele is then transmitted to *Z*’s offspring with a probability of 1/2 for each, and thus gamete *a* has a probability of 1/4, and *b*, *c* each have a probability of 1/8 of carrying the mutant allele. Different coefficients of identity are calculated depending on whether the probability of gametes carrying the allele are defined conditional on the allele frequencies in some particular generation.

**Table 1.** Symbols and their meaning

| Symbol | Meaning |
| --- | --- |
| $\Delta_n$ | One of Jacquard's (1970) nine condensed modes of identity by descent, described in table 2. |
| $\Delta$ | Without a subscript, delta is used to symbolize the coefficient of cofraternity. |
| $E[a]$ | The expectation, or mean, of the random variable $a$. |
| $\hat{a}$ | An estimate of $E[a]$ |
| $\mu_n(a)$ | The $n^{th}$ central moment of $a$, defined as $E[(a - E[a])^n]$. |
| $\mu(a, b, \dots)$ | A mixed central moment of $a$ and $b$ and omitted variables, defined as $E[(a - E[a])(b - E[b])\dots]$. |
| $\gamma_{\tilde{X}, Y}$ | The inbred-relatedness, which is the probability of sampling a locus where individual $X$ is inbred and also related to individual $Y$. |
| $\Theta_{X, Y}$ | The coefficient of co-ancestry, or kinship, between individuals $X$ and $Y$, which is the probability that an allele in individual $X$ is identical by descent to an allele in individual $Y$. Also denoted by $\theta$ and $\Phi$ in some texts. |
| $\Delta_{\tilde{X}, \tilde{Y}}$ | The second-order zygotity correlation of individuals $X$ and $Y$, which is the probability that there are two pairs of alleles which are each identical by descent. Either both individuals are inbred but unrelated, or the individuals are genetically identical but not inbred. |
| $\delta_{\tilde{X}, \tilde{Y}}$ | The fourth-order zygotity correlation of genotypes between individuals $X$ and $Y$, which is the probability that all four alleles in the two individuals are identical by descent. |
| $\epsilon_{XY}$ | The allele frequency estimation error, the difference in allele frequencies between individuals $X$ and $Y$. |
| **Quantitative-genetic terms** |  |
| $\sigma_A^2$ | The additive genetic variation of the population. |
| $\sigma_D^2$ | The dominance genetic variation of the population. |
| $\sigma_{ADI}$ | The covariance of additive and dominance effects in inbred individuals. |
| $\sigma_{DI}^2$ | The variance of dominance effects in inbred individuals. |

### Calculating P(a) and P(b)

Unfortunately, the sampling of a haplotype is a singular event. We cannot directly measure the probability of sampling an allele in a particular haplotype in a particular individual. However, we can use the distribution of genotypes within the population as a whole, in combination with some statistical model, as an estimate of that probability. The simplest statistical model we can use is a uniform distribution, where the presence or absence of an allele is independent and identically distributed among all haplotypes; in this case our estimate of *P*(*a*) is simply the frequency of the allele in the population. This is not an unreasonable procedure, but we need to keep in mind that *P*(*a*) is being calculated on the condition that the frequency of a is *ƒ*(*a*) = *θ* in the population, and would be more properly written as *P*(*a*|*ƒ*(*a*) = *θ*), and not simply *P*(*a*).

### Conditional independence

By conditioning on the current allele frequency in a panmictic population, *ƒ*(*a*) = *θ*, we can theoretically remove the genotypic correlation created by genetic drift. If an individual *X* gives no information about the genotype of individual *Y*, aside from aiding in the estimation of the allele frequency in the population, then the genotypes of *X* and *Y* can be made independent by conditioning on that allele frequency. This process implicitly occurs when molecular markers are used to estimate genotypic correlation coefficients, because allele frequencies must be measured in the current generation. Pedigree based methods neglect this step by assuming that allele frequencies are known *a priori*, and as a result the effects of drift are not removed explicitly or implicitly. As a result genotypic correlation coefficients increase monotonically as pedigrees become more extensive (Speed and Balding 2015).

### Negative correlations

Conditioning on the allele frequency in the current generation will not in general make the genotypes of all distantly related individuals independent. Individuals in a population have varying degrees of relatedness, and different allele frequencies will be necessary to make different pairs of individuals independent. No allele frequency will make all sufficiently distant individuals independent. For instance, in Figure 1 an allele that originates in individual *Z* has an uncon-ditional probability *P*(*X*)= 3/16 and *P*(*Y*)= 1/16 ^3^ of being sampled in individuals *X*, *Y*, respectively. The unconditional probabilities of sampling the allele are independent because the allele descends through entirely separate lineages. Because *X* and *Y* share no ancestors that possess the mutant allele except for *Z*, we do not learn anything about the frequency of the mutant allele in *Y*’s ancestors from *X*. Yet if we condition on the allele frequency in the parental generation (from which gametes *a*, *b*, *c* and *d* were sampled), then individuals *X* and *Y* become negatively correlated.

It is easiest to see how this negative correlation arises if we first think of conditioning on the generation containing *X* and *Y* themselves. For instance, consider the case where we condition on sampling a single copy of the mutant allele among the four alleles of *X* and *Y*. In this case learning that the allele was sampled in *Y* tells us with certainty that the allele was not sampled in *X*, because we already know the allele was sampled only once. A similar, though less severe, process occurs when we condition on the parental generation. Learning that individual *Y* possesses an allele makes it more likely that copies of that allele were present in *Y*’s parents, and if the total number of alleles in the parental generation is known, then it becomes less likely that *X*’s parents had copies of the allele, and *vice versa*. A negative genotypic correlation indicates that an allele tends to be sampled from either individual *X* or individual *Y*, but usually not both.

Negative correlations can occur in large unstructured populations. Stochastic difference in reproductive success occur between families delineated by any level of relatedness (e.g. a family at the level of first cousins, all second cousins, etc.), and just as in the preceding example, larger families will contribute more alleles to estimates of allele frequencies than small families. As a result conditioning on allele frequency estimates will create negative correlations between some pairs of individuals. These negative correlations are a fundamental aspects of population structure which describe these differences in reproductive success, and they do not become trivial in large populations. Negative correlations are unfamiliar in the context of population structure because of the tradition of envisioning these correlations as estimates of probabilities, but later we will observe negative genotypic correlation coefficients among individuals in real populations, so we will need to have some understand of the mechanism that creates them.

### Expression for pairwise genotypic correlation

Two random variables (for example *a* and *b* in Figure 1) that can each be in one of two possible states (*a* = 0 and *a* = 1) can take on four states jointly (*a* = 1 and *b* = 0, *b* = 1 and *a* = 0, etc.), each with their own associated probability. These four outcomes have three degrees of freedom (one degree of freedom is lost because the four states sum to 1), and thus we need three parameters to describe the overall distribution: the probability that *a* = 1, *P*(*a*), the probability that *b* = 1, *P*(*b*), and some parameter that describes the association of *a* and *b*. An obvious choice for this association term is the correlation coefficient between a and b, which is a covariance that has been normalized by the geometric mean of the standard deviations of the univariate distributions, and can be written as:

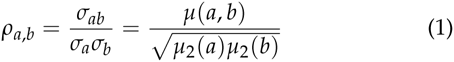

where *σ*_ab_ and *μ*(*a*, *b*) are two different notations for the covariance between *a* and *b*, and *σ*_a_^2^ and *μ*(*a*) are two different notations for the variance of *a*. In general *μ*_n_(*a*) is the *n*^*th*^ central moment of *a* and is defined as *E*[(*a* — *E*[*a*])^n^], where *E* denotes the expectation, or raw moment, of a variable.

The covariance is related to the correlation coefficient by 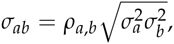 so we can write the probability of *a* and *b* in terms of the means *P*(*a*) and *P*(*b*), and the correlation coefficient *ρ*_*a,b*_ as:

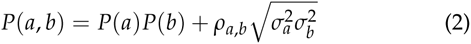

By substituting *a*’ = 1 — *a* for *a*, we can write the probabilities of *P*(*a*’*b*), *P*(*ab*’) and *P*(*a*’*b*’) in a similar manner, *e.g.*:

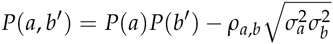

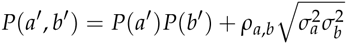

If we ignore the particular genealogy shown in Figure 1, and instead consider the general case, the two gametes composing individual *X* and the two gametes composing individual *Y* could have been sampled from different populations, and the frequencies of alleles could differ in these populations, so it may be the case that *P*(*a*) ≠ *P*(*b*) ≠ *P*(*c*) ≠ *P*(*d*), where *P*(*a*) and *P*(*b*) are the allele frequencies in *X*’s parents and *P*(*c*) and *P*(*d*) are the frequencies in *Y*’s parents. The correlation coefficient *ρ*_*a,b*_ can still be used to describe the genotypic correlation between *a* and *b*, and the joint probabilities can be written in the form of eq. 2, though determining the allele frequencies may be difficult.

There are six unique ways to choose two items from four items if items can be chosen only once, and the order of choice does not matter (*i.e.* four choose two is six). As a result there are six ‘second-moment’correlation coefficients between any four random variables. These are just correlation coefficients in the ordinary sense, but need to be distinguished from the higher-moment coefficients describing the statistical association of three or four random variables, which are introduced later. Two of these six ‘second-moment’ coefficients are inbreeding coefficients, which are the correlation coefficients of the gametes that fused to make an individual:

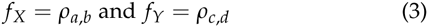

The other four second-moment coefficients are generally not considered separately; instead, their arithmetic average defines the coefficient of coancestry:

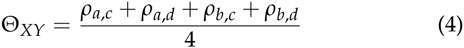

The use of the arithmetic average is not an arbitrary choice. If the two diploid individuals *X* and *Y* produce two gametes *e* and *ƒ*, the gametes will have a second-moment correlation coefficient of:

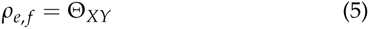

There are sixteen possible paired genotypes of two individuals at biallic loci (*P*(*abcd*), *P*(*a’bcd*), *P*(*a’b’cd*), etc.), and in order to fully describe the joint probability of these sixteen paired genotypes several parameters are needed. Co-skewness and co-kurtosis coefficients both arise naturally when describing three-and four-variable statistical associations, in the same way that covariance arises when describing two-variable associations. While we will not use the co-skewness or co-kurtosis themselves, the third and fourth-moment correlation coefficients are related to co-skewness and co-kurtosis and share properties with them.

Skewness (rather than co-skewness) measures the asymmetry of a probability distribution and is defined as *μ*_3_ · *μ*^−3/2^; it measures whether observations have a tendency to be either larger (for positive skewness) or smaller (for negative skewness) than the mean. Co-skewness is the multivariate analog of skewness that represents the tendency of jointly distributed variables to simultaneously take on values on the same (for positive) or different (for negative) side of the means (*i.e.*, major-allele frequencies) of the distribution, and is defined as *μ*(*a,b,c*) · (*μ*_2_(*a*)*μ*_2_(*b*)*μ*_2_(*c*))^−1/2^. While co-skewness isadimen-sionless parameter—because the numerator and denominator are the same order—it does not estimate genotypic correlations. The co-skewness of a haplotype with itself is simply the skewness, whereas the genotypic correlation of a haplotype with itself is one.

To obtain a statistic that does not vary as a function of allele frequency, and thus estimates genotypic correlations, we normalize the third central mixed moment by the third moments of the univariate distributions, yielding:

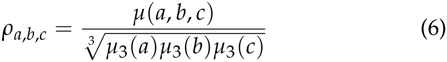

This notation is adapted to emphasize the similar form and behavior of this parameter to a correlation coefficient, and we will refer to *ρ*_*a,b,c*_ as the third-moment correlation coefficient. (The choice of the geometric mean of central moments is described in more detail in the Supplemental section SC). The third-moment correlation can be used in a fashion similar to the second-moment correlation coefficient to write probabilities of the joint distribution of three variables, *e.g.*:

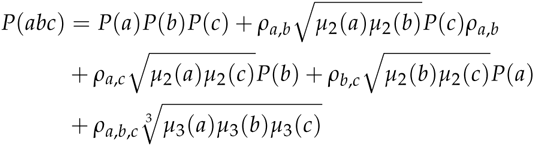

A total of four third-moment correlations exist between four haploid genomes, which are grouped into two arithmetic averages:

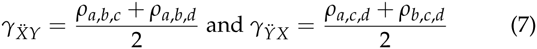

We call these terms the inbred-relatedness coefficients, because they describe the probability of sampling a site where the first index individual (*X* for *ϒ*_*ẌY*_ or *Y* for *ϒ*_*ΫX*_) is inbred and related to the second indexed individual (through either one or both of the alleles in the second indexed individual). The symbols *ϒ*_*ẌY*_ and *ϒ*_*ΫX*_ are adopted from Cockerham (1971). Again, the arithmetic mean is not an arbitrary choice, but instead represents a formulation that allows us to express the third-moment correlation of gametes produced by individuals.

The final term is the fourth-moment correlation coefficient. This coefficient estimates the faction of sites There are three modes of identity by descent (Δ_1_, Δ_2_ and *Delta*_7_ in Figure 2) where the zygosity of the two individuals is guaranteed to be identical, is defined as:

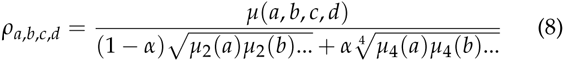

where *μ*_2_(*a*), *μ*_2_(*b*)…are the second central moments, *μ*_4_(*a*), *μ*_4_(*b*)…are the fourth central moments, the dots denote the omission of the moments of *c* and *d*, and *α* is a term that describes the fraction of the fourth-moment correlation coefficient that arises from the fourth-moment component as described bellow. Unlike the second-moment and third-moment coefficients, where all identity modes contributing to the genotypic correlation describing the same basic kind of relationship (either two alleles that are IBD or three alleles that are IBD), the fourth-moment coefficient is composed of two different kinds of relationships: a relationship where all four alleles are IBD, and a relationship of two pairs of two alleles, either in the same (Δ_2_) or different (Δ_7_) individuals, which are each IBD. These are the second-and fourth-moment components of *ρ*_*a,b,c,d*_: Δ_*ẌΫ*_ and *δ*_*ẌΫ*_ and can be expressed in terms of *ρ*_*a,b,c,d*_ as:

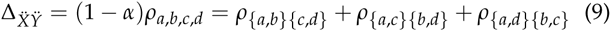

and

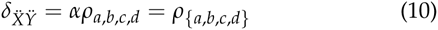

**Figure 2.**
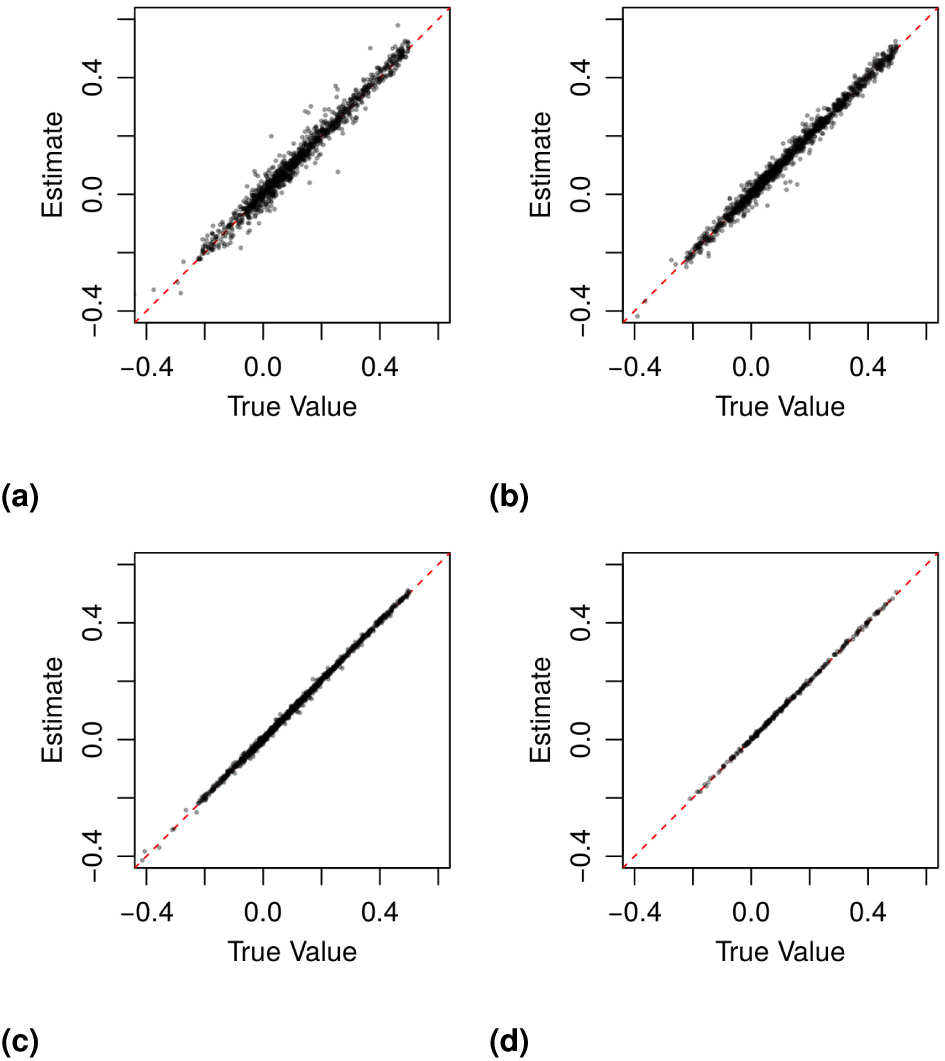
Results from 10,000 simulation using 3× (a) and 10× (b) coverage at 5,000 loci; and 3× (c) and 10× (d) coverage at 100,000 loci; allele frequencies were drawn from a triangular distribution as described and reported without error. All seven genotypic correlation coefficients are graphed jointly. A summary of biases and mean squared error can be found in Table S3.

The notation {*a, b*}{*c, d*} in the subscript specifies which groups of alleles are IBD. This notation becomes necessary when there are several different ways in which the relationship can be partitioned. The coefficient Δ_*ẌΫ*_ is the sum of the terms Δ_*Ẍ+Ϋ*_ and Δ_*Ẍ · Ϋ*_ used by Cockerham (Holland et al. 2003) and becomes the the coefficient of cofraternity Δ, in the absence of inbreeding. The term *δ*_*ẌΫ*_ is Jacquard’s (1970) Δ_1_ and is also used by Cockerham (1971). We call *ρ*_*a,b,c,d*_ the zygosity correlation coefficient because it describes how zygosity *(i.e.*whether an individual is heterozygous or homozygous) is correlated between individuals, with Δ_*ẌΫ*_ called the second-moment zygosity correlation component and *δ*_*ẌΫ*_ called the fourth-moment zygosity correlation component because they depend on the second-and fourth-moments of the univariate distributions respectively.

Every set of nine probabilities for jointly sampling the genotypes of individuals *X* and *Y* can be transformed into a unique set of eight parameters. Two of these parameters are the probabilities of sampling the minor allele in individuals *X* and *Y*, and the other six are the genotypic correlation coefficients relating *X* and *Y*—*ƒ*_*X*_, *ƒ*_*Y*_, Θ_*XY*_, *ϒ*_*ẌY*_, *ϒ*_*ΫX*_, and *μ*(*a, b, c,d*). However the joint genotypic probabilities are only defined for some values of these eight parameters (*i.e.* the relationship is not a one-to-one correspondence). Strong correlation of the genotypes of the individuals places a constraint on the difference of the probability of sampling the minor allele in both individuals. If correlations are close to one then the difference of minor allele frequencies must be close to zero, but when correlations are close to negative one the difference must be close to one minus the minor allele frequency.

Finally, it is the joint central moment *μ*(*a, b, c,d*) which is specified by a set of nine genotypic probabilities, and not a genotypic correlation coefficient, either Δ_*ẌΫ*_, *δ*_*ẌΫ*_ or *ρ*_*a,b,c,d*_. The values of Additionally the coefficients Δ_*ẌΫ*_ and *δ*_*ẌΫ*_ require a range of allele frequencies to estimate, because they describe how the denominator of Eq. 8 changes as a function of allele frequencies. The relationship of these coefficients to the nine condensed modes of IBD is shown in Table 2.

**Table 2.**
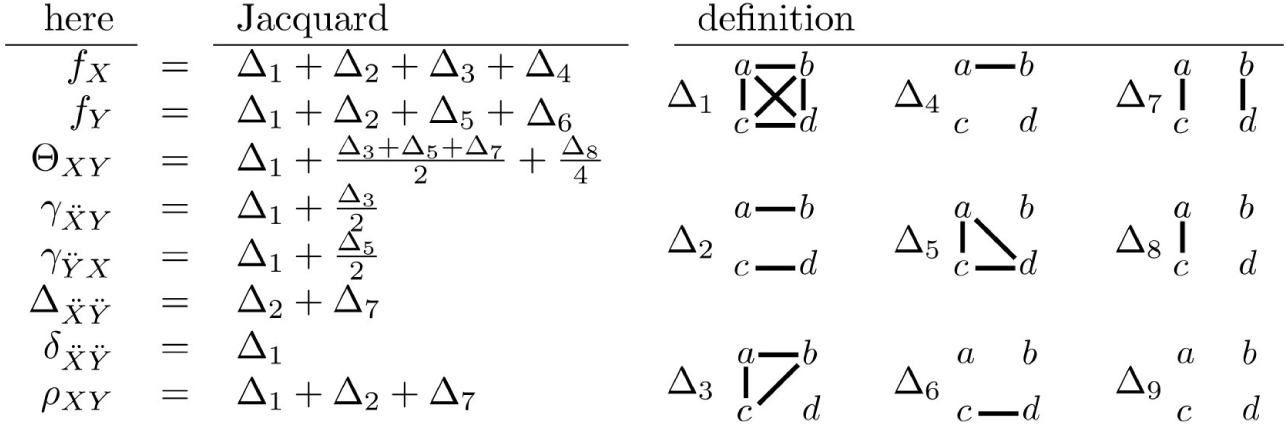
Relationship between genotypic correlation coefficients and modes of identity by descent (IBD) On the right are the nine identity modes relating two individuals. Alleles *a* and *b* belong to individual *X*, and alleles *c* and *d* belong to individual *Y*. Alleles that are IBD are connected by solid lines. On the left, theses nine modes can be used to obtain the coefficients of co-ancestry (Θ), inbreeding (*ƒ*), and cofraternity (Δ), along with the coefficients that we introduce: the inbred relatedness (*γ*), identity (*δ*), and zygosity (*ρ*).

Three condensed IBD modes can be expressed in terms of these seven coefficients: Δ_1_ = *δ*_*ẌΫ*_, Δ_3_ = *ϒ*_*ẌY*_ — *δ*_*ẌΫ*_ and Δ_5_ = *ϒ*_*ΫX*_ - *δ*_*ẌΫ*_. Although we do not consider these coefficients for three or more alleles in this paper, if more alleles were present Δ_*ẌΫ*_ could be separated into its components Δ_2_ and Δ_7_, and the remaining six condensed IBD modes could be estimated from linear combinations of the genotypic correlation coefficients presented here.

### A complete model of genetic covariance in populations

These genotypic correlation coefficients will be important in the analysis of quantitative traits. In the absence of inbreeding and epistasis, the genetic covariance of quantitative traits can be defined as:

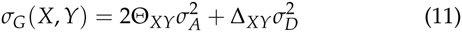

In the presence of inbreeding, but absence of epistasis, this becomes:

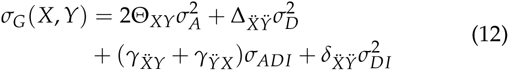

where *σ*_ADI_ is the covariance of additive and dominance effects in inbred individuals (2*D*_1_ in Cockerham), and σ^2^_DI_ is the variance of dominance effects in inbred individuals (*D*_2_^*^ in Cockerham) (Cockerham 1983, Abney et al. 2000, see Lynch and Walsh 1998). In the absence of inbreeding the terms *ϒ*_*ẌY*_, *ϒ*_*ΫX*_, *δ*_*ẌΫ*_ are all 0, and Δ_*ẌΫ*_ = Δ_*XY*_, so Eq 12 reduces to Eq. 11. However, even in ostensibly outbred populations, particular individuals will have small but statistically significant amounts of inbreeding (or outbreeding) because real populations are not perfectly panmictic. As a result, estimates of genetic covariance using Eq. 12 should be more accurate than estimates which assume a perfectly panmictic population and use Eq. 11. As with the additive and dominance genetic variance, care should be taken in the verbal interpretation of the covariance of additive and dominance effects in inbred individuals (*σ*_ADI_) and the variance of dominance effects in inbred individuals (σ^2^_DI_), as these parameters describe properties of populations and do not describe modes of gene action.

### Estimating IBD coefficients from population-genomic data

There are three steps in computing the coefficients of relatedness using sequence data. First, an estimate of the allele frequencies in the populations from which the two individuals are sampled must be obtained; second, the genotypes of the two individuals being compared must be estimated; and third, a relatedness estimate must be constructed from this information. The first two steps are closely related, because they both involve making inferences about genotypes from sequence data, and we use a framework that jointly estimates sequence error rates and allele frequencies at each site (Maruki and Lynch 2015), implemented in the program mapgd (Ackerman et al. In prep). A benefit of this approach is that it not only produces unbiased estimates of population parameters when depth of coverage is low, but because we explicitly model sequencing as two discrete events (the random sampling of chromosomes from an individual followed by the random distribution of errors among reads), we can assess whether the observed data are consistent with our statistical model. The likelihood equation used to estimate the parameters, Eq. S13, can be transformed into an cumulative distribution function describing the probability of obtaining data of lower likelihood than the observed data, Eq. S16. By limiting the analysis to genomic sites consistent with the model, we can remove sites that potentially suffer from sequencing or assembly artifacts (Supplemental Section S4).

The third and final step in the process is to estimate the genotypic correlation coefficients from genotypic probabilities and allele frequencies. We do this by maximizing a likelihood equation that describes the probability of observing the pattern of reads given a set of genotypic correlation coefficients. For a particular site, the likelihood equation is the product of the three terms 1) the probability *P*(*G*_*x*_ = *i*|*X*_*k*_) that individual *X* has genotype *G*_*x*_ = *i* given that the ‘quartet’ (*i.e.* counts of A,C,G and **T’**s observed at the site) ***X***_*k*_ is observed at site *k*; 2) the corresponding probability *P*(*G*_*y*_ = *i*|*Y*_*k*_) for individual *Y*; and 3) the probability *P*(*G*_*X*_ = *i*, *G*_*Y*_ = *j*|***θ***) of observing the pair of genotypes *G*_*X*_ and *G*_*Y*_ in the two individuals given a set of genotypic correlation coefficients ***θ***. When estimating genotypic correlation coefficients the terms *P*(*G*_*x*_ = *i*|*X*_*k*_) and *P*(*G*_*y*_ = *i*|*Y*_*k*_) are simplified from Eq. 4b in Lynch (2008), because the error rate and major-and minor-allele identities have been estimated prior to this calculation by mapgd from the population data. The term *P*(*G*_*X*_ = *i*, *G*_*Y*_ = *j*|***θ***) is taken from the joint genotypic distributions developed in the previous section (defining the coefficients of identity by descent) with ***θ*** being the vector of the seven genotypic correlation coefficients. Explicit forms of *P*(*G*_*X*_ = *i*, *G*_*Y*_ = *j*|***θ***) and *P*(*G*_*X*_ = *i*|***X***_k_) in terms of allele frequencies, genotypic correlation coefficents, error rates and nucleotide quartets are given in Supplemental Section S2.

Because the genotypes (of which there are three: 1. homozygous for the major-allele, 2. heterozygous, and 3. homozygous for the minor-allele) are mutually exclusive events, we sum across them (*i,j* ∈ {1,2,3}), and assume that observations at the *n* different loci are independent. The product (or the sum of the logs) of the likelihood of the data at each site gives us the likelihood give us the likelihood of the data overall:

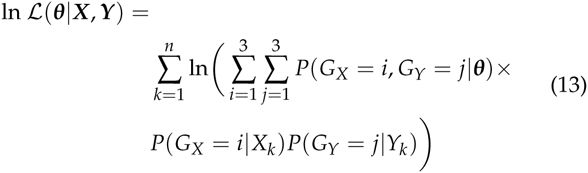

where ***X***_*k*_ and ***Y***_*k*_ are the quartets of individuals *X* and *Y*, at site *k* respectively.

The maximum likelihood estimates of the genotypic correlation coefficients are found by maximizing Eq. 13. This is done with a numerical method, currently the sequential least-squares programming method implemented in the python SciPy module. Genotypic correlation with large magnitude limit the possible difference of allele frequencies between individuals. Our model assumes that this difference is zero. Unfortunately this causes *P*(*G*_*X*_ = *i*, *G*_*Y*_ = *j*|***θ***) to become undefined when negative correlations are large and minor allele frequencies are small. While it may be possible to address this problem in a rigorous manner, currently we arbitrarily set *P*(*G*_*X*_ = *i*, *G*_*Y*_ = *j*|***θ***) to 0 when it is undefined and normalize the remaining probabilities. Because our model behaves poorly when minor-allele frequencies are low, we recommend ignoring sites with minor-allele frequencies less than 0.05.

### Simulations

Two kinds of simulations were performed to test our methods. The first simulation was designed to examine the statistical performance of mapgd’s estimation procedure. For these simulations the genotypes of two individuals were simulated and the genotypic score…allele frequency in the population from which these individuals were sampled was either be reported without error or reported with simulated sampling error. Related individuals were generated by selecting 1 to 7 genotypic correlation coefficients, which were then assigned a random value between −1 and 1. The remaining coefficients assigned a value of 0. The coefficients were checked to ensure that the joint probability distribution was defined for minor-allele frequencies between 0.1 and 0.4, and a file was generated with either 5 × 10^4^,10^5^ or 10^6^ SNPs at either 3× or 10 × coverage. Allele frequencies were drawn from a triangular distribution with mean mean 0.1, minimum 0 and maximum 1. Finally binomially-distributed noise representing the sampling error in estimates of allele frequencies was introduced for sampling 10,100 and 1000 individuals.

We also simulated a genomics study by creating 150 bp reads that were aligned to a simulated reference genome using bwa (Li and Durbin 2010). Allele frequencies within the reference population followed a Pareto-distribution and were estimated from the sequence of 96 unrelated individuals, because under a neutral expectation allele frequencies are Pareto-distributed with *α* = 1/2*N*.

The accuracy of our estimates of relatedness and inbreeding were compared to the programs vcftools, king and plink. Unfortunately, no other method currently exists that estimates the coefficients *ϒ* or *δ* but, several these programs can calculate the coancestry (Θ) and cofraternity (Δ) coefficients in the absence of inbreeding, denoting them as either *k*_1_ and *k*_2_ after Cotter-man (1940), or *IBD1* and *IBD2* after Suarez et al. (1978). In the absence of inbreeding these terms are equivalent to Θ and Δ respectively, and are usually verbally described as representing the probability that one (*k*_1_ or IBD1) or two (*k*_2_ or IBD2) alleles are identical by descent between a pair of individuals. For details on the operation of mapgd see Ackerman *et al.* (In prep); for vcftools, see Yang et al. (2010); for king, see Manichaikul et al. (2010); and for plink, see Purcell et al. (2007).

## Results

### Simulation Results

Our maximum-likelihood estimation procedure produces accurate and precise estimates of all seven relatedness components, even when depth of coverage is minimal (Figure 2). The bias (the expected error—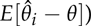 and the mean squared error (MSE) 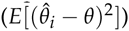 depend on the number of loci sampled, the depth of coverage, and the accuracy of allele-frequency estimates. Individuals in typical metazoan populations may have 1 × 10^6^ and 10×10^6^ informative SNPs^4^ and even at 3× coverage we can estimate all seven components with MSE less than 2.4 × 10^−5^. When the number of loci used is reduced to 100,000 and 5,000, the largest MSE increases to 1.5 × 10^−4^ and 28.4 × 10^−4^, respectively.

The most biased estimators are the inbreeding coefficients *ƒ*_*X*_ and *ƒ*_*Y*_, which overestimate the amount of inbreeding by 0.003 when coverage is low and only 5,000 SNPs are used. This bias arises largely from the small number of SNPs used in the analysis and not from low coverage, with a similar bais occurring at 10× coverage.

Errors in the estimation of allele frequencies upwardly bias estimates of *ƒ*_*X*_, *ƒ*_*Y*_, Θ_*XY*_, *ϒ*_*ẌY*_, *ϒ*_*ΫX*_ and *δ*_*ẌΫ*_ and downwardly bias Δ_*ẌΫ*_. This bias is roughly independent of depth of coverage and number of loci used, but does depend on the number of individuals sampled, resulting in an upward bias of 0.01 to *ƒ*_*X*_, *ƒ*_*Y*_, Θ_*XY*_, *ϒ*_*ẌY*_, and *ϒ*_*ΫX*_ and *δ*_*ẌΫ*_ when 50 individuals are used to estimate allele frequencies.

The errors of five estimates of genotypic correlation coefficients (*ƒ*_*X*_, *ƒ*_*Y*_, Θ_*XY*_, *ϒ*_*ẌY*_ and *ϒ*_*ΫX*_) are uncorrelated, whereas the errors of the estimates of the two zygosity coefficients (Δ_*ẌΫ*_ and *δ*_*ẌΫ*_) have a strong negative correlation to each other (*r*^2^ = 0.67), and consequently these two terms also have the largest MSEs. However, when 10^4^ or more loci are used, the MSE of both Δ_*ẌΫ*_ and *δ*_*ẌΫ*_ are < 1.5 × 10^−4^.

We compared the performance of mapgd to vcftools (Yang et al. 2010), which can estimate Θ (using the-relatedness option) and f (using the-het option), and to king (Manichaikul et al. 2010) and plink (Purcell et al. 2007), which both estimate Θ and Δ, on our mock genomics study of 98 individuals. Using the default settings of each program, we find that the maximum-likelihood method implemented in mapgd substantially reduces the bias and MSE of identity coefficients compared to vcftools, king, and plink, particularly when genotyping error rates are high. At a coverage of 3×, king underestimates coancestry by 48%, plink by 49%, and vcftools by 27% for outbred siblings. Increasing the coverage to 10× reduces the bias to 8%, 8% and 5%, respectively, and all of the programs are essentially unbiased at 30× coverage (Table 3). However, unlike the method we present here, high coverage does not ensure accurate estimates from vcftools, plink, or king, because they are all sensitive to assumptions regarding inbreeding to various degrees. The program king seems to be particularly sensitive to these assumptions, generally estimating that inbred siblings are unrelated (*i.e.* Θ = 0).

**Table 3.** Bias (×10^3^) and MSE (×10^4^) of mapgd, vcftools, king, and plink in estimating coancestry (Θ), cofraternity (Δ_*ẌΫ*_), and inbreeding (*ƒ*_*X*_) for outbred and inbred siblings. The values of all 7 IBD coefficients are listed in table S5. Results are from 100 simulations on a 400 kbp genome containing ~10,000 SNPs. See supp:simulations for SNP filtering parameters.

| Coancestry | | outbred ($\Theta = \frac{1}{4}$) | | inbred ($\Theta = \frac{1}{2}$) | |
| --- | --- | --- | --- | --- | --- |
| Program | Cov. | Bias | MSE | Bias | MSE |
| mapgd | $3\times$ | -10. | 2.2 | -11.6 | 2.5 |
| vcftools | $3\times$ | -68 | 50 | -150 | 230 |
| king | $3\times$ | -120 | 170 | -450 | 2000 |
| mapgd | $10\times$ | -6.9 | 0.73 | -8.2 | 1.5 |
| vcftools | $10\times$ | -12.0 | 1.8 | -16 | 3.3 |
| king | $10\times$ | -21 | 4.7 | -520 | 2700 |
| mapgd | $30\times$ | -6.4 | 0.69 | -7.2 | 1.0 |
| vcftools | $30\times$ | -5.7 | 0.7 | -4.5 | 1.0 |
| king | $30\times$ | -3.1 | 0.29 | -500 | 2500 |
| Cofraternity | | outbred ($\Delta = \frac{1}{4}$) | | inbred ($\Delta = \frac{3}{8}$) | |
| Program | Cov. | Bias | MSE | Bias | MSE |
| mapgd | $3\times$ | -7.2 | 22. | -0.3 | 23 |
| plink | $3\times$ | -20 | 7.1 | 72 | 52 |
| king | $3\times$ | 370 | 1400 | 230 | 520 |
| mapgd | $10\times$ | -1.7 | 4.3 | -1.4 | 5.8 |
| plink | $10\times$ | 11 | 2.3 | 110 | 120 |
| king | $10\times$ | 58 | 36 | 400. | 1700 |
| mapgd | $30\times$ | 1.2 | 4.3 | 8.6 | 6.1 |
| plink | $30\times$ | 1.7 | 1.6 | 120 | 150 |
| king | $30\times$ | -3.5 | 1.7 | 360 | 1300 |
| Inbreeding | | outbred ($f_X = 0$) | | inbred ($f_X = \frac{1}{2}$) | |
| Program | Cov. | Bias | MSE | Bias | MSE |
| mapgd | $3\times$ | -14 | 7.1 | -33 | 18 |
| vcftools | $3\times$ | 99 | 100. | -100. | 110 |
| plink | $3\times$ | 72 | 54 | -150 | 220 |
| mapgd | $10\times$ | -8.5 | 1.6 | -7.0 | 1.3 |
| vcftools | $10\times$ | 15 | 3.5 | -11 | 9.5 |
| plink | $10\times$ | 12 | 2.4 | -7.4 | 1.2 |
| mapgd | $30\times$ | -9.0 | 1.9 | -7.6 | 1.6 |
| vcftools | $30\times$ | -0.1 | 1.9 | -3.8 | 1.3 |
| plink | $30\times$ | -2.2 | 1.4 | -2.4 | 0.9 |

In contrast to the poor performance of vcftools, plink and king on low coverage sequence or with relatives with complex relationships, mapgd gives accurate and unbiased estimates across all simulated coverage and relationships (Table 3). This robust estimation comes with a substantial computational cost, with estimates from mapgd taking longer than the other methods. The major computational hurdle for accurate estimation of relatedness is the accurate calculation of allele frequencies. But, this investment in computational time results in a substantial increase in the accuracy of allele-frequency estimation (Figure 3).

**Figure 3.**
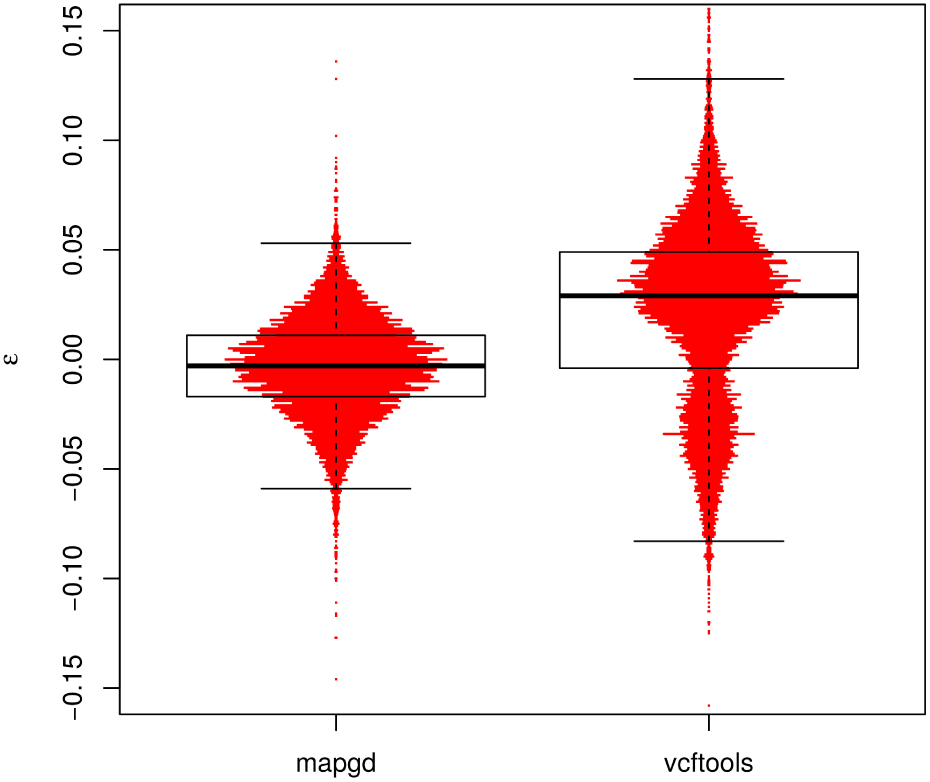
Violin plot of the errors (*ε*) of allele-frequency estimates from the programs mapgd and vcftools. The horizontal width of the red bars represent the frequency of observations with the corresponding values of *ε*. The black box shows the median (heavy black line), boundaries of the upper and lower quartile (so that 50% of all errors are contained within the box) and the whiskers denote observations within 1.5 interquartile range of the upper and lower quartiles. Results from ~ 10,000 estimates of a population of 98 individuals with 3× coverage. Allele are drawn from a neutral spectrum. Allele frequencies in vcftools are calculated by the vcftools-freq command.

The surprisingly poor performance of king in the presence of inbreeding arises from an attempt by the program to compensate for population structure; While this may be successful under other circumstances, here it infers that the single inbred pair of siblings are a unique sub-population. Disabling this option, with the–homo argument, substantially reduces the bias of king’s coancestry calculations, but it still compares unfavorably with mapgd.

### Analysis of Daphnia population genomic data

*Daphnia pulex* is a microcrustacean commonly found in ephemeral ponds. During much of the year *Daphnia* produce re-produce asexually, but many *Daphnia* can produce resting eggs, called ephippia, through sexual reproduction (sexuals). Since only resting eggs survive the winter in ephemeral ponds, sexual *Daphnia pulex* must have sex at least once a year in order to persist. Some *pulex* can only produce resting eggs asexually (asexuals) allowing them to persist between years without sex (Hebert and Crease 1980).

Samples of 96 *Daphnia pulex* were collected from four ephemeral ponds: Kickapond, Portland Arch, Busey, and Spring Pond South (See Fig S3 for a map of locations). Early season samples were collected to minimize the chance of sampling clone mates (genetically identical individuals), which are produced asexually by all female *Daphnia* at one to four week intervals but cannot survive the winter. Each of these samples was screened using 6 allozyme loci for evidence of a shared multilocus genotype as a rudimentary screen for asexuals. Because no population appeared to have asexuals in high abundance, all of these populations were sequenced to ~15× average coverage on an Illumina MySeq. The reads were aligned to a reference genome (Colbourne et al. 2011) and analyzed with mapgd (Supplemental Section S6).

Three of the four *Daphnia* populations showed mild but significant outbreeding (Figure 4). The population displaying inbreeding (Spring Pond South) also contained a number of clone mates (12 genetically distinct individuals sampled 74 times). Genetically identical individuals were also isolated from Portland Arch (7 genetically distinct individuals sampled 17 times), but not from the two other populations. We analyzed these individuals for asexual markers described in previous studies (Tucker *et al.* 2013, Xu *et al.* 2015), and found that three of the twelve groups of clone mates in Spring Pond possessed asexual makers, giving a total of 24 putative asexual individuals. The putata-tive asexuals were outbred compared to the local population (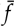 = —0.10 ± 0.02 vs. 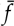 = 0.03 ± 0.01). We removed the two chromosomes known to have a hybrid origin in asexual individuals (Xu et al. 2013 Tucker *et al.* 2013, Xu *et al.* 2015) from the analysis (Figure S1a), but individuals with asexual markers still appear outbred compared to the local population.

**Figure 4.**
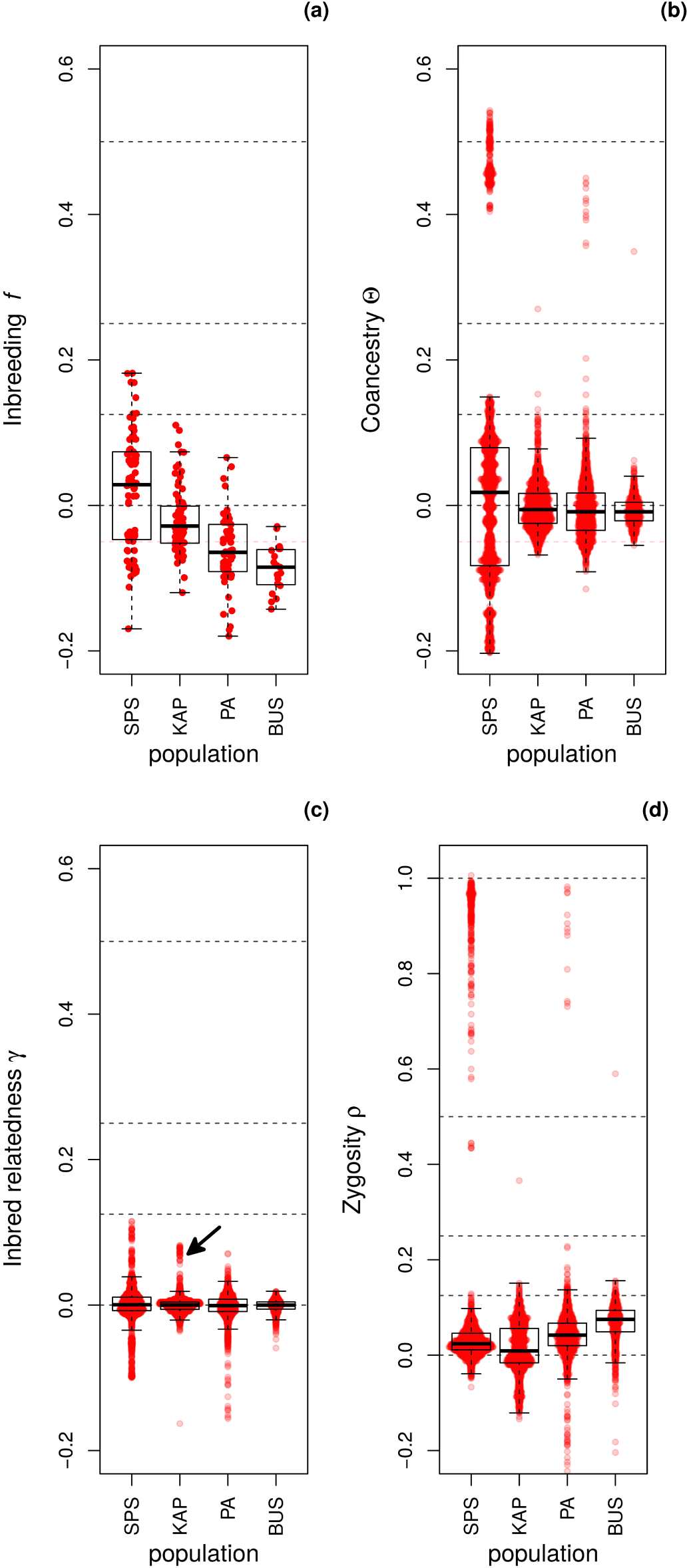
Box plots of the genotypic correlation coefficients of inbreeding—*ƒ*(a), coancestry—Θ(b), inbred relatedness— *ϒ*(c), and zygosity—*ρ*_*ẌΫ*_ (d) estimated in the four Daphnia populations. The arrow in figure (c) indicates a group of 9 individuals analyzed in more detail in Figure S1b. The individual pairwise estimates are shown as red points. Because Θ, *ϒ*, and *ρ* are pairwise estimators, ~4,500 comparisons exist in each population for these coefficients, while only ~95 estimates exist for *ƒ*. Dashed lines are placed at 1, 0.5, 0.25, 0.125, and 0 to allow easier assignment of relationship status

In addition to groups of clone mates displaying Θ ≈ 0.5 and Δ ≈ 1.0^5^ (Figure 4b and 4d), several other general patterns are apparent. Strongly negative coancestry values separate the individuals with asexual markers from other individuals in Spring Pond South (figure 4b). These individuals are also separated by negative inbred relatedness values (Figure 4c).

A group of nine individuals in Kickapond displayed elevated inbred relatedness values (figure 4c, arrow). These individuals had elevated inbreeding, coancestry, and second and fourth-order zygosity correlations with each other group, and a generally negative coancestry with the rest of the population (Figure S1b).

The second and fourth order zygosity correlation components show a strong negative correlation with each other (figure S2), which is consistent with the behavior of estimation error in our simulations (S3), and also consistent with the behavior of the estimators when there is population structure (data not shown). Because of these negative correlations, the zygosity correlation coefficient (*ρ* = Δ_*ẌΫ*_ + *δ*_*ẌΫ*_) was used in analyses.

Kickapond and Busey ponds had no clonally mates, but Portland Arch and Busey both had a pair of first-order relationships (Θ ≈ 0.25, Table 4). The coancestry (Θ) of BUS-109 and BUS-110 is consistent with these two *Daphnia* being full siblings, and this is supported by their relatively large cofraternity coefficient (Δ), but was some what less than expected for full siblings (Δ=0.5). However, BUS-10 has a large inbreeding value and the relationship with BUS-11 is inconsistent with some form of inbred half-siblings because the inbred relatedness values are too low. It may be that the low coverage (2.5×) of BUS-10 caused an overestimate of inbreeding and coancestry values, and an underestimate of confraternity, although this error would have to be more severe than errors seen in simulation at 3× coverage. PA-12 and PA-108 also demonstrate a relatively high coancestry, but too low to be full siblings, and they have a low cofraternity as well. In this case it may be that PA-12 and PA-108 are half-siblings, but that population structure is obscuring their relationship. Two pairs of individuals in Kickapond, one pair in Busey and one pair in Portland Arch are consistent with half-siblings with varying degrees of inbreeding (table 4). The coancestry statistics are all ~0.125 and other genotypic correlation coefficients are small. However, most of these relationships are still confounded with population structure, as demonstrated by the generally negative inbreeding coefficients. There are also half-sibling-like relationships in Spring Pond South, however, the performance of the estimators is very sensitive to estimations of allele frequencies, and the allele-frequency calculations in Spring Pond South are based on the smallest number of individuals due to the the large number of clone mates sampled. Because estimation of the genotypic correlation coefficients depends on accurate allele frequency estimates, significant bias may exist in these estimates.

**Table 4.** The values of genotypic correlation coefficients from the 6 individuals from Kickapond, Portland Arch, and Busey displaying coancestry of half-sib ships or higher (Θ > 1/8), excluding clone mates. Spring Pond South is excluded due to small sample size. Genotypic correlation coefficients significantly different from zero (*χ*^2^ > 10.7, corrected for the 42 comparisons in the table) are indicated with a dagger (^†^).

| Clone X | Clone Y | $f_X$ | $f_Y$ | $\Theta_{XY}$ | $\gamma_{\bar{X}\bar{Y}}$ | $\gamma_{\bar{Y}\bar{X}}$ | $\delta_{\bar{X}\bar{Y}}$ | $\Delta_{\bar{X}\bar{Y}}$ |
| --- | --- | --- | --- | --- | --- | --- | --- | --- |
| BUS-10 | BUS-11 | 0.14$^\dagger$ | -0.03$^\dagger$ | 0.34$^\dagger$ | 0.08$^\dagger$ | 0.01$^\dagger$ | 0.00 | 0.42$^\dagger$ |
| PA-12 | PA-108 | -0.07$^\dagger$ | -0.00 | 0.20$^\dagger$ | -0.02$^\dagger$ | 0.00 | -0.01$^\dagger$ | 0.08$^\dagger$ |
| PA-024 | PA-051 | -0.06$^\dagger$ | -0.08$^\dagger$ | 0.13$^\dagger$ | 0.00 | -0.00 | 0.01 | 0.05$^\dagger$ |
| PA-112 | PA-04 | -0.04$^\dagger$ | -0.08$^\dagger$ | 0.13$^\dagger$ | -0.02$^\dagger$ | -0.02$^\dagger$ | -0.00 | 0.03 |
| KAP-048 | KAP-099 | -0.00 | 0.03$^\dagger$ | 0.15$^\dagger$ | -0.01$^\dagger$ | 0.01$^\dagger$ | -0.00 | 0.01 |
| KAP-113 | KAP-120 | -0.04$^\dagger$ | -0.03$^\dagger$ | 0.13$^\dagger$ | -0.01$^\dagger$ | -0.01$^\dagger$ | 0.00 | 0.00 |

## Discussion

Here we reemphasize the importance of allowing genotypic correlation coefficients to take on negative values—not only because doing so decreases the bias of estimates, but also because the expected correlations can indeed be negative. The framework outlined here is the first method to provide accurate estimates of zygosity correlation coefficients in the presence of inbreeding, and also the first method to provide estimates of inbred-relatedness coefficients from population-genomic data. This method provides accurate and nearly unbiased estimates on very low coverage data. The genotypic correlation coefficients recovered from the four populations of *Daphnia* provide insight into population structure by recovering close family relationships and separating distinct subpopulations. Finally, because we use a maximum likelihood method, log-likelihood ratio tests can be used to evaluate alternative models for statistical significance.

There is a long history of interpreting IBD coefficients as correlation or regression coefficients, and it has long been recognized that these coefficients can be extended to include coefficients relating an arbitrary number of individuals (Wright 1922, Cockerham 1971). For instance, Wright’s *FST* can be thought of as the genotypic correlation of all members of a defined subpopulation to each other. However, it has generally been thought that such extensions would prove impractical to calculate, or that the number of coefficients would increase very quickly^6^. But by grouping partitions of the same order, as we do with the second-and fourth-order zygosity correlation coefficients, the number of coefficients can be substantially reduced. Programs capable of algebraic manipulation may make it possible to extend this method to groups larger than pairs of individuals.

Methods of recovering individual specific estimates of allele frequencies need to be developed. This phrase may seem self-contradictory since an individual does not in an ordinary sense of the word have an allele frequency. But an individual does have a probability of possessing an allele, and as discussed in the section **Calculating *P*(*a*) and *P*(*b*)**, this probability is usually taken to be identical with the allele frequency in the population. Because population structure can cause this frequency to differ for different groups of individuals, and these groups do not need to be discrete, every individual can have a unique probability of possessing an allele, and thus, in some sense, can have an individually unique allele frequency. While Eqs 3 through 9 formally allow differences in ancestral allele frequencies between individuals, we have not made an attempt to estimate this parameter. However, even with these limitations, the ability to detect and characterize complex relationships in wild populations, such as that between PA-12 and PA-108, should be a boon to researchers. While many programs can detect some relationships in panmictic populations, mapgd is unique in that its estimates are accurate in the presence of inbreeding, and it provides additional genotypic correlation coefficients not estimated by other programs.

The draft genome to which reads were aligned in this study is known to suffer from a number of artifacts, particularly “allelic splits”, where the two alleles of a gene assemble as paralogous genes in the reference, and “paralog collapse” where paralogous genes are assembled at a single locus (Denton et al. 2014). While the goodness of fit statistic was developed in part to detect and remove these artifacts, it’s performance has not been carefully analyzed in this paper, and it is possible that artifacts undetected by the goodness of fit statistic influence our estimates. The inbreeding of an individual is the coancestry of the parents, and while slightly negative coancestry is common in these populations, there are few individuals with Θ < −0.05, but many individuals with *ƒ* < −0.05 (Figure 4b and Figure 4a). This may be artifactual. High coverage increases the power of the goodness of fit test, and should increase our ability to discern artifacts. While an elevation of inbreeding is seen in low coverage individuals, inbreeding estimates appear stable once coverage is greater than 5×, implying that mapgd’s estimates are relatively robust once sequencing depth is reasonable. Nevertheless, the robustness of these estimates needs to be reevaluated when a better reference genome becomes available.

In this study, genotypic correlations provided insight into a number of aspects of population structure. The three separate groups of asexuals in Spring Pond South could be distinguished from sexuals by their large negative inbreeding, coancestry, and inbred relatedness values. This result is consistent with limited or no crossing between the asexuals and the sexuals within the pond. A group of nine individuals in Kickapond with a coancesty of Θ ≈ 0.1 form a clear subpopulation (Figures 4c and S1b), although we have not explored what factors drive this structure.

An average of two close relatives were found in the three Daphnia populations (excluding clone mates), which may seem surprisingly high given that only 96 individuals were sampled. However, ~4,500 comparisons of relatedness were made between individuals in each population, so the possibility of obtaining related individuals is much greater than naive intuition would suggest. Our ability to find closely related individuals in random samples highlights the potential power of a genomic based approach in wild populations.

One strength of the maximum likelihood framework is that it allows the assessment of significance of relationships through a log-likelihood ratio test. Virtually all of the pairs of *Daphnia* had highly significant departures from an unrelated status (*i.e. p* ≪ 10^−4^ after multiple correction), in stark contrast to the behavior of our estimators in simulations when the true value of the parameters are known to be zero. In addition to geographic structure created by the isolation of populations from each other, the dormancy of resting eggs can create temporal structure. This temporal and geographic structure may be responsible for many of the significant departures from panmixia. In this case the log-likelihood ratio test is, at least in a sense, working correctly. Since most populations may have some form of substructure owing to variation in family size and demography *etc*., it may be desirable to find a method that considers some aspects of population structure as part of the null model.

While much work remains to be done, our estimators already have excellent performance when coverage is minimal. The ability to use low coverage data for population-genomic studies will greatly reduce the cost of these studies. Even if the additional parameters estimated by methods outlined here are unneeded, the ability to recover accurate and precise estimates of coancestory, inbreeding, and cofraternity estimates on low coverage sequence may be of general use. We hope that these properties, and others discussed in the text, will make the general coefficients of genotypic correlation useful to the research community.

## Acknowledgments

We would like to thank colleagues for their helpful comments on this paper, especially Lydia Bright; Thanks to W. Kelley Thomas and the Hubbard Center for Genome Studies for their continued help with the sequencing; The Center for Genomics and Bioinformatics and especially James Ford for the preparation of libraries; Kimberly Young of the *Daphnia* Growth Facility and her army of undergraduates for their tireless work maintaining lines; Ben Fulton and others at the Pervasive Technology Institute for help with mapgd. This research was made possible by a grant from the National Science Foundation to support the development of methods for the analysis of population genomic data (DEB – Award 1257806) and a grant from the National Institute of Health to support population genomics research in *Daphnia pulex* (GM101672-01A1).

## Footnotes

1 46 chromosome, 2 crossover events per chromosome per meiosis and 2 meiotic events

2 Each of the 184 crossover events has a 50% chance of being resolved in the same direction in both siblings, so the 95% CI interval

3 These are the probabilities that a gamete generated by *X* or *Y* will carry the mutant allele. Half of the time a gamete generated by *X* will inherit the allele carried by gamete *a*, which has the mutant allele with probability 1/4, and half of the time *X*’s new gamete will inherit the allele carried by gamete *b*, which has the mutant allele with probability 1/8, so *P*(*X*)= 3/16. A similar calculation is done for *Y*.

4 Roughly genome size times heterozygosity. For humans this is 3 × 10^9^ × 10^−3^ = 3 × 10^6^ and in *Drosophila* 2 × 10^8^ × 3^−2^ = 6 × 10^6^

5 This is the expectation for clone mates because their zygosity state is always identical (Δ ~ 1.0) and alleles are only identical when comparing paternal to paternal, or maternal to maternal alleles, which occurs half of the time (Θ ~ 0.5).

6 The number of partitions of a set of size *n* are denoted by the *n*^*th*^ Bell number which is defined by the recursive relationship 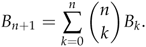 These numbers become large very quickly For 9 items there are *B*_9_ = 21,147 partitions and for 10 items there are *B*_10_ = 115,975 partitions.

